# Slow conduction and spatial dispersion of repolarization are intrinsic properties of cardiomyocyte electrophysiology that contribute to proarrhythmia in an iPSC model of hypertrophic cardiomyopathy

**DOI:** 10.1101/2023.07.20.549952

**Authors:** Seakcheng Lim, Melissa M. Mangala, Mira Holliday, Henrietta Cserne Szappanos, Samantha B. Ross, Whitney Liang, Ginell N. Ranpura, Jamie I Vandenberg, Christopher Semsarian, Adam P. Hill, Livia C. Hool

## Abstract

Hypertrophic cardiomyopathy (HCM) is an inherited heart muscle disease; characterised by left ventricular wall thickening, cardiomyocyte disarray, and fibrosis, and is associated with arrhythmias, heart failure and sudden death. However, it is unclear to what extent the electrophysiological disturbances that lead to sudden death occur secondary to the structural changes in the myocardium, or as a result of intrinsic properties of the HCM cardiomyocyte. In this study, we used an induced pluripotent stem cell model of the Arg403Gln variant in myosin heavy chain 7 (*MYH7*) to study ‘tissue level’ electrophysiological properties of HCM cardiomyocytes. For the first time, we show significant slowing of conduction velocity and an increase in local spatial dispersion of repolarisation - both well-established substrates for arrhythmia - in monolayers of HCM cardiomyocytes. Analysis of rhythmonome protein expression in R403Q cardiomyocytes revealed dramatically reduced connexin-43, sodium channels, and inward rectifier channels – a three-way hit that combines to reduce electrotonic coupling between HCM cardiomyocytes and slow cardiac conduction. Our data therefore represent a novel, biophysical basis for arrhythmia in HCM, that is intrinsic to cardiomyocyte electrophysiology. Later in the progression of the disease, these proarrhythmic electrical phenotypes may be accentuated by fibrosis and myocyte disarray to contribute to sudden death in HCM patients.

## 1. Introduction

Hypertrophic cardiomyopathy (HCM) is an autosomal-dominant inherited cardiac disorder with a prevalence of up to 1 in 200 that can result in arrhythmias, heart failure and sudden death^1,2^. Disease-causing variants associated with HCM most commonly occur in sarcomere genes including *MYH7* (myosin heavy chain), *MYBPC3* (myosin binding protein C3), and *TNNT2* (troponin T), with variants in the *MYBPC3* gene being the most common cause^3^. HCM is characterised clinically by left ventricular hypertrophy ≥ 15 mm in the absence of loading conditions^4,5^ and is also associated with fibrosis, myocyte disarray and altered energy metabolism^6^. Electrical conduction delays and dispersion of repolarisation, both risk predictors for arrhythmias due to an increased risk of re-entry^7^, are also clinical features of HCM^8^. However, the correlation between these conduction defects and the histopathology of HCM is limited^9–11^. As a result, the mechanisms underlying electrical dysfunction in HCM, and its role in sudden death in patients is unclear.

The missense arginine to glutamine substitution at position 403 in *MYH7* (R403Q) is a HCM variant that causes severe disease, characterised by early-onset and progressive myocardial dysfunction, with a high incidence of sudden cardiac death^12,13^. In mice, homozygous αMHC^403/403^ results in neonatal lethality, while the heterozygous mouse (αMHC^403/+^) also demonstrates cardiac dysfunction, myocyte disarray, hypertrophy and fibrosis^14^. However, consistent with some observations in patient cohorts, neither the extent nor the location of fibrosis correlated with electrical mapping of conduction properties in this mouse model^10^. Furthermore these histopathological changes also did not correlate with the propensity for arrhythmia. It has also been reported that tachyarrhythmias are observed at a far earlier age than the onset of hypertrophy^11^, and that dysfunction of cardiac calcium current and alteration of mitochondrial metabolism occurs prior to the onset of myopathy^13^. Each of these pieces of evidence suggests an alternative pathway for arrhythmic substrate formation in HCM, such as electrical remodelling, that is at least partly independent of alterations in the structure of the myocardium.

In this study, we used a patient-derived iPSC model of the *MYH7* R403Q mutation (*MYH7*^403/+^))^15^ to investigate whether there are intrinsic electrical properties of HCM cardiomyocyte electrophysiology that provide a biophysical basis for arrhythmia in the absence of structural alteration of the myocardium. Our data show for the first time that in *MYH7*^403/+^ cardiomyocytes, a dramatic reduction in expression of connexin-43 and sodium channel proteins results in reduced conduction velocity. Accompanying this is an increase in spatial dispersion of repolarisation that establishes potential proarrhythmic substrates and may provide a biophysical basis that contributes to sudden arrhythmic death in HCM patients.

## 2. Methods

### 2.1. Generation of patient-specific human induced pluripotent stem cells and general cell culture maintenance

Human induced pluripotent stem cells (iPSCs) were derived from a patient carrying the HCM causing mutation p.Arg403Gln in *MYH7*. Peripheral blood mononuclear cells (PBMCs) were isolated and reprogrammed iPSCs as previously described^15^. Patient informed consent, use and generation of patient-derived iPSCs complied with national guidelines with oversight by the Sydney Local Health District Committee (Protocol X19-0108 and ETH00461). iPSC colonies were maintained in a defined, feeder cell-free medium, mTeSR1 PLUS™ (Stem Cell Technologies, Tullamarine, AUS) and the extracellular matrix, Matrigel hESC-qualified matrix (Corning Inc., NY, USA), and passaged as aggregates using ReLeSR™ passaging reagent (Stem Cell Technologies). Brightfield images were captured on the Zeiss Primo Vert inverted microscope and processed on the Zeiss ZEN Lite 3.4 software (Zeiss, Oberkochen, DEU).

### 2.2. CRISPR-Cas9 gene editing of iPSCs

The guide RNA, below, for generation of isogenic control lines were designed using an online tool (crispr.mit.edu) and ordered as oligos to be cloned into pSpCas9(BB)-2A-Puro (PX459) V2.0 (Addgene, MA, USA).

Sequence 5’ to 3’: CATTGCCCACTTTCACCTGA

Homology directed repair (HDR) donor oligos were designed to flank the Cas9 cut site and ordered as ultramers that contained 2 phosphorothioate bonds on each end of the oligo (Integrated DNA Technologies, IA, United States).

Donor template for correcting MYH7 Arg403Gln:

C*C*TCATGGGGCTGAACTCAGCCGACCTGCTCAAGGGGCTGTGCCATCCTCGGG TGAAAGTGGGCAATGAGTACGTCACCAAGGGGCAGAATGTCCAGC*A*G

iPSCs were plated as single cells and transfected 24 h later with 1 µg plasmid and 5 µL 100 µM donor oligo using LipofectamineTM Stem (Thermo Fisher, MA, USA) according to manufacturer’s protocol. After transfecting for 24 h, cells were selected with 0.5 µg/mL puromycin for a further 24 h. Single cells were selected into a 96-well plate for Sanger sequencing to determine successfully edited clones which were further expanded into established cell lines. Off target analysis involved Sanger sequencing of ten potential guide RNA off-target sites (http://www.rgenome.net/cas-offinder/), Sanger sequencing of all TP53 exons and molecular karyotyping (Victorian Clinical Genetics Services, VIC, AUS) to ensure genomic integrity.

### 2.3. Cardiac differentiation

Cultures of iPSCs at 70-80 % confluence were dissociated into a single cell suspension with TryPLE™ (Thermo Fisher) for 7 minutes at 37 °C, 5 % CO_2_. Cells were then plated between 450,000 - 750,000 cells per well of a Matrigel coated 12-well tissue culture plate in mTeSR1 PLUS™ (Stem Cell Technologies) supplemented with 10 µM ROCK inhibitor (Y-27632) (Reprocell, Beltsville, USA) and differentiated into cardiomyocytes using the StemDiff Cardiomyocyte differentiation kit (Stem Cell Technologies) as per manufacturers protocol.

### 2.4. Multi-electrode array (MEA) recordings

At day 15 (± 2 days) beating iPSC-CMs were dissociated using a two-step protocol described previously (Mills et al., 2016). Briefly, iPSC-CMs were incubated with 0.2 % Collagenase Type I (Thermo Fisher), in PBS supplemented with 20 % fetal bovine serum, for 45 minutes at 37 °C, 5 % CO_2_ then centrifuged 300 x *g* for 3 minutes. iPSC-CMs were incubated in 0.25 % Trypsin-EDTA (Thermo Fisher) for 10 minutes at room temperature, then filtered through a 40 µm cell strainer. iPSC-CMs were plated at a density of 10,000 cells/well of an Axion Biosystems E-Stim+ Classic MEA 48-plate (Axion Biosystems, GA, USA), and field potentials/conduction velocities were recorded at day 30 using the Maestro-APEX MEA system (Axion Biosystems). For field potential durations, repolarisation time was measured from the point of maximum slope of the depolarization spike to the peak of the repolarizing ‘t wave’. Spontaneous and paced activities of iPSC-CMs were recorded at 37 °C, 5 % CO_2_. Acquisition and analysis were performed using AxIS v2.5.1.10 software (Axion Biosystems), The Cardiac Analysis Tool (Axion Biosystems), in house MEA analysis software (Victor Chang Cardiac Research Institute, NSW, Australia), and MATLAB R2021a (MathWorks, MA, USA).

### 2.5. Western blotting

At day 45 (± 2 days) protein was extracted from beating iPSC-CMs in protein lysis buffer (1 X Pierce™ RIPA buffer (Thermo Fisher); 1 X PhosSTOP cocktail (Sigma, MO, USA), 1 X cOmplete protease inhibitor (Sigma). Protein concentration was quantified using the Bradford assay using BSA as a standard. 25 µg of total protein was loaded into precast 10 % BIO-RAD Mini-PROTEAN® TGX Stain-FreeTM SDS-polyacrylamide gel (Bio-RAD Laboratories), then electrophoretically transferred to 0.2 µm nitrocellulose membrane (Trans-Blot® TurboTM Transfer Pack, Bio-RAD Laboratories) using the Bio-RAD Trans-Blot® TurboTM Transfer System. Blots were probed with the following primary antibodies: rabbit polyclonal anti-KV1.4 (Alomone, #APC-167, 1:500), rabbit polyclonal anti-KV1.5 (Alomone, #APC-004, 1:500), guinea pig polyclonal anti-KV2.1 (Alomone, #AGP-109, 1:1000), rabbit polyclonal anti-KV4.2 (Alomone, #APC-023, 1:500), guinea pig polyclonal anti-Kir2.1 (Alomone, #AGP-044, 1:500), rabbit polyclonal anti-K2P3.1 (TASK1) (Alomone, #APC-024, 1:500), rabbit polyclonal anti-Kir6.2 (Alomone, # APC-020, 1:500), rabbit polyclonal anti-KV11.1 (HERG) (Alomone, #APC-109-F, 1:500), rabbit polyclonal anti-KCNE1 (IsK, MinK) (Alomone, #APC-163, 1:500), rabbit polyclonal anti-KCNQ1 (KV7.1) (Alomone, #APC-168, 1:500), rabbit polyclonal anti-SAP97 (ThermoFisher, #PA1-741, 1:1000), rabbit polyclonal anti-NaV1.5 (Alomone, #ASC-005, 1:500), rabbit polyclonal anti-CaV1.2 (Alomone, #ACC-003, 1:500), rabbit polyclonal anti-connexin 43 (Cell Signalling, #3512, 1:1000), rabbit monoclonal anti-GAPDH (Cell Signalling, #2118, 1:2000) and subsequently with the pre-absorbed, polyclonal goat anti-rabbit IgG (H & L) HRP (Abcam, #ab97040, 1:10000), or polyclonal goat anti-guinea pig IgG (H & L) HRP (Abcam, #ab97155, 1:10000) secondary antibody. Blots used for re-probing with multiple antibodies were stripped using stripping buffer, consisting of (in mM): Tris HCl pH 6.8: 62.5, 2-mercaptoethanol: 100, 2 % SDS, 30 min at 50 °C. Densitometry was performed for each antibody using ImageJ software^16^. Background subtracted intensity values were normalized to loading control GAPDH signal intensity detected on the same blot. All immunoblot experiments were run as triplicate as minimum.

### 2.6. Nanostring analysis

RNA was harvested with 100 µL of QIAzol Lysis Reagent (Qiagen, Hilden, DE) per well of an MEA plate. RNA was pooled from 3-4 wells of an MEA plate and then extracted using the miRNeasy kit (Qiagen). RNA levels were measured using the nanoString nCounter® PlexSet™ (nanoString, WA, USA) according to the manufacturer’s instructions and gene expression analysed using nSolver software (nanoString). A two-stage Benjamini, Krieger & Yekutieli procedure was implemented for controlling the false discovery rate (FDR)^17^. A FDR of 5% and a fold-change of 2 in gene expression were considered significant.

### 2.7. Data analysis and statistical tests

Data analysis, plotting and statistical tests were performed using Microsoft Excel (Microsoft Office 2016, Microsoft, WA, USA), and GraphPad Prism v9 (GraphPad Software, CA, USA). All data are expressed as mean ± SEM unless otherwise indicated. Data was analysed using a mixed effect model using the *glmfit* function in Matlab, with individual differentiations as fixed categorical values (N) and technical replicates (n) as random variables. Estimated marginal means were derived from the generalised linear mixed effect models using the *emmeans* package^18^, with comparison between differentiations undertaken using a Wald test on input contrasts. In all cases, significance was determined with *p*-values < 0.05. Violin SuperPlots of data were created using Violin SuperPlots in Matlab^19^, error bars on violin plots represent estimated marginal means from the generalised linear model and their standard errors. Raw data and code for statistical analysis and data visualisation are available for download from Zenodo (web address).

## 3. Results

### 3.1. Generation of iPSC lines for MYH7^403/+^ and isogenic control

An iPSC line was reprogrammed from a 38-year-old female HCM patient (II:1, Figure 1A) with the pathogenic myosin heavy chain 7 (*MYH7*) p.Arg403Gln mutation in myosin heavy chain 7 (*MYH7*^403/+^) as previously described^15^ (includes quality control assessment of karyotype, pluripotency and trilineage differentiation capacity). For this study, we generated an isogenic control line by correcting the pathogenic variant in *MYH7* using CRISPR-Cas9 genome editing (*MYH7*-C^+/+^) (Figure 1B). DNA sequencing confirmed the A>G correction at c.1280G>A (Figure 1C) while no off-target edits were detected at the top 10 predicted sites of p53 identified using PCR (Supplementary Table 1). *MYH7*^+/+^ iPSC lines successfully generated colonies of iPSCs with classical morphology of tightly packed cells with a high cell-to-nucleus ratio (Figure 1D). In all subsequent experiments, this CRISPR-corrected line was used as the comparator for functional and molecular assessments.

**Figure 1.**
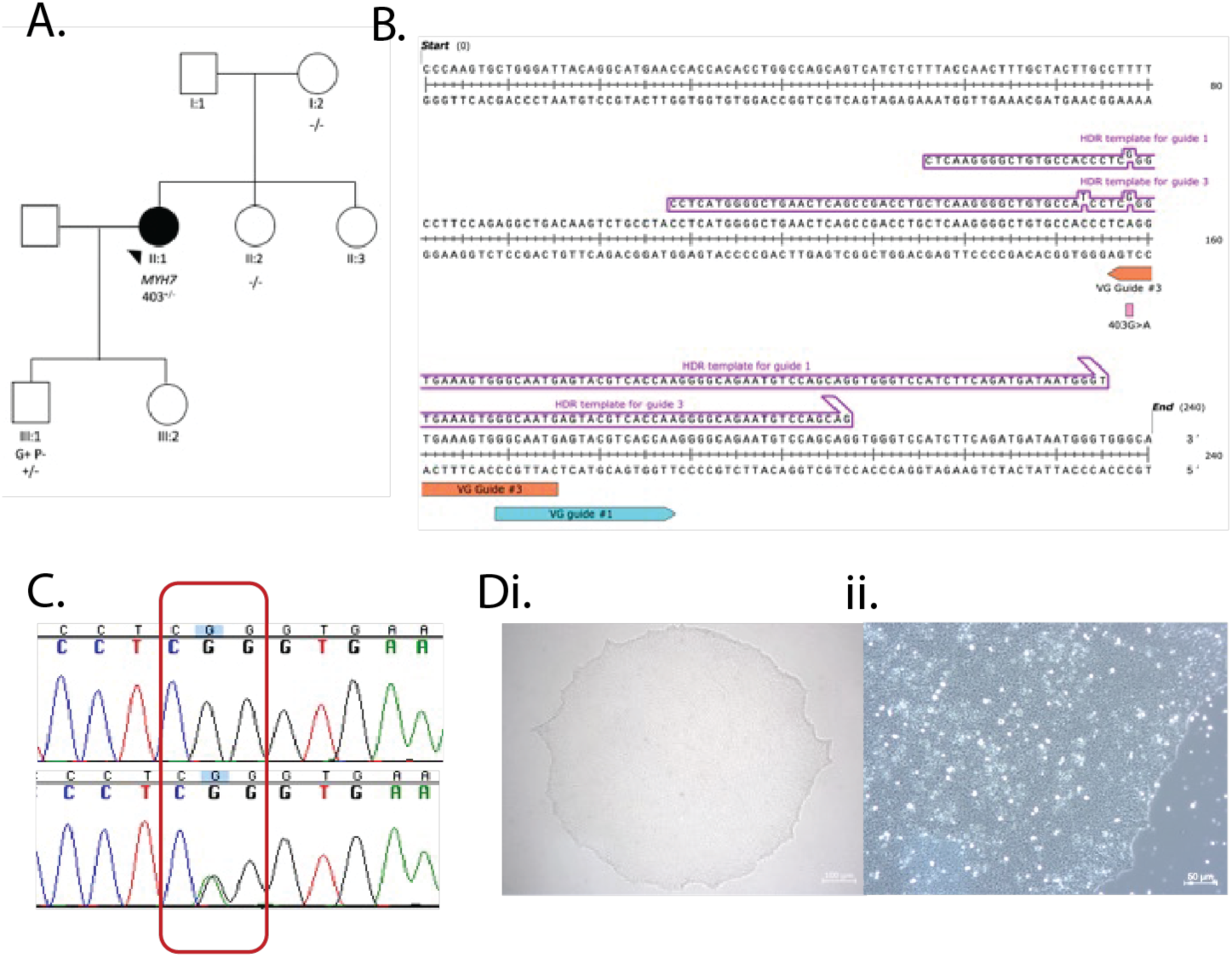
Generation of isogenic control iPSC line for MYH7^403/+^. A) MYH7^403/+^ iPSC were reprogrammed from a HCM patient (II:1) with the MYH7 p.Arg403Gln variant as previously described^14^. B) HDR template used to CRISPR-correct MYH7 p.Arg403Gln variant in iPSCs. C) DNA sequencing showing correction of (p.Arg403Gln c.1208G>A). D) Brightfield images of CRISPR-corrected iPSCs at 4 x and 10 x magnification.

### 3.2. MYH7^403^*^/+^* modifies electrophysiology of iPSC-derived cardiomyocytes

Cardiomyocytes derived from iPSC for both *MYH7*^403/+^ and *MYH7*-C*^+^*^/+^ were seeded onto microelectrode arrays and extracellular field potentials – an *in vitro* surrogate of the electrocardiogram (ECG) - were recorded from spontaneously beating monolayers of cells (Figure 2A). The R403Q variant affected both depolarisation and repolarisation properties of the cardiomyocytes. Specifically, *MYH7*^403/+^ showed a reduction in the slope of the depolarisation spike of the field potential (Figure 2B), a measure of the propagating action potential upstroke, from −0.14 ± 0.02 V/s in *MYH7*-C^+/+^ to −0.04 ± 0.02 V/s in *MYH7*^403/+^, and increased the Fridericia corrected field potential duration (FPDc; Figure 2C) from 281.0 ± 21.6 ms in *MYH7^+^*^/+^ to 318 ± 21.5 ms, in *MYH7*^403/+^, reflecting slowed repolarisation of the cardiac myocytes (consistent with our previous studies showing prolonged action potential duration in *MYH7*^403/+^ cardiomyocytes^20^). In addition, *MYH7*^403/+^ cardiomyocytes displayed more irregular/arrhythmic beating (Figure 3), significantly increasing the coefficient of variation of beat rate from 7.0 ± 5.2 % to 17.8 ± 5.2 % (Figure 3C). For *MYH7*^403/+^, a variety of arrhythmic phenotypes were observed, ranging from more subtle beat-to-beat variability in cycle length (Figure 3A/Bii) to the presence of ectopic depolarisations, often tightly coupled to regular/sinus beats (Figure 3A/Biii).

**Figure 2.**
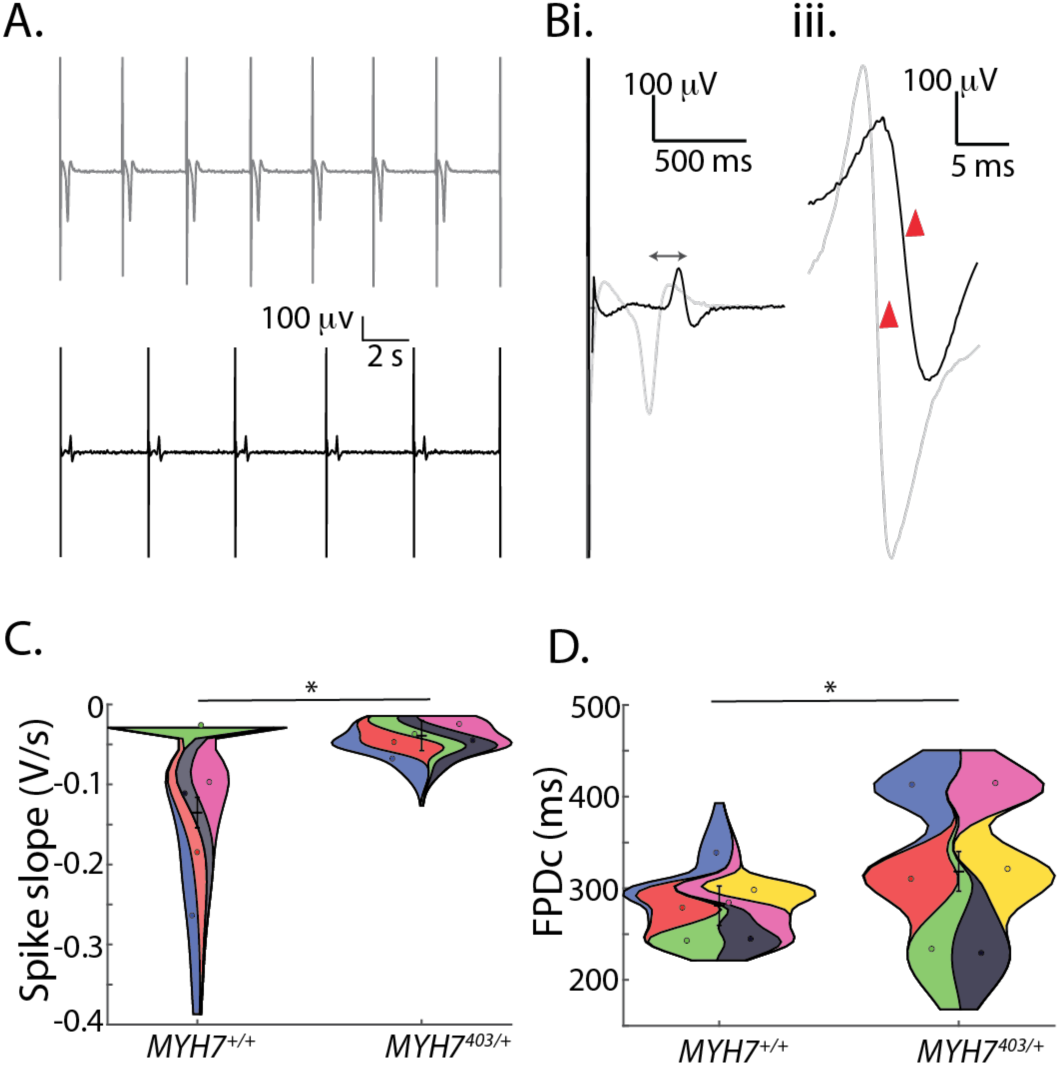
*MYH7*^403/+^has altered depolarisation and repolarisation in iPSe-eMs. A) Representative field potentials from *MYH7*^+/+^ (grey, top) and *MYH7*^403/+^ ipse-eMs (black, bottom). B) Zoomed view highlighting differences in repolarization time (field potential duration; (i)) and slope of the depolarization complex (red arrows; (ii)). e) Violin plot summarizing depolarization slope for *MYH7*^+/+^ (n = 73 replicates/wells from N = 6 differentiations) and *MYH7*^403/+^ (n = 89 replicates/wells from N = 6 differentiations) showing significantly slower depolarization for *MYH7*^403/+^. (N = 6, n = 66 wells). E) Violin plot showing rate corrected field potential duration (FPDc) for *MYH7*^+/+^ (n = 85 replicates/wells from N = 6 differentiations) and *MYH7*^403/+^ (n = 111 replicates/wells from N = 6 differentiations) showing significantly prolonged repolarisation for *MYH7*^403/+^.

**Figure 3.**
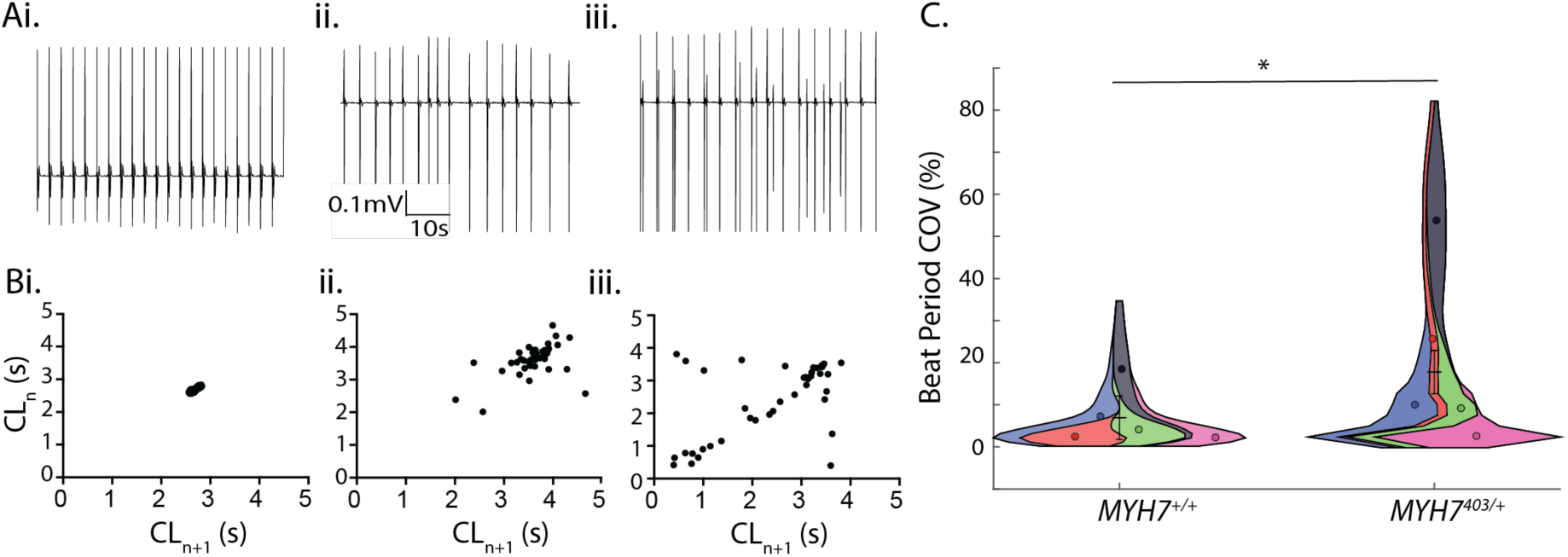
MYH7^403/+^ causes irregular/arrhythmic beating in iPSe-eMs. A) Example traces of field potentials from MYH7^+/+^ (i) and MYH7^403/+^ (ii-iii). B) Poincare plots summarising beat rate irregularity from 2 minute recordings in A. e) Violin plot summarising beat period irregularity for MYH7^+/+^ (n = 88 replicates/wells from N = 6 differentiations) and MYH7^403/+^ (n = 111 replicates/wells from N = 6 differentiations) showing a significant increase in beat period irregularity for MYH7^403/+^.

### 3.4. Reduced electrical coupling in MYH7 R403Q slows conduction and induces spatial dispersion of repolarisation

A reduced slope of the depolarisation spike (Figure 2C) might reflect multiple electrophysiological phenomena, including cellular factors such as a reduced density of sodium currents, or macroscopic ‘tissue-level’ properties such as altered cell-cell coupling leading to slowed propagation of the activating wavefront. We therefore next measured conduction velocity in monolayers of iPSC cardiomyocytes. Conduction maps for *MYH7*-C^+/+^ and *MYH7*^403/+^ iPSC-CMs, coloured according to activation time measured at individual electrodes across the microelectrode array, are shown in Figure 4A. In these examples, the activation wavefront takes longer to propagate from bottom right to top left of the array in *MYH7*^403/+^ compared to *MYH7*-C^+/+^. Overall, propagation time was increased from 13 ± 1 ms to 25 ± 1 ms respectively (Figure 4B), reflecting slower conduction velocity. This slowed conduction velocity smight suggest that syncytia of *MYH7*^403/+^ cardiomyocytes are less tightly coupled than *MYH7*^+/+^. In normal hearts, the level of electric coupling between cells attenuates differences in electrical properties between individual cells [20]. In the context of reduced electrical coupling in *MYH7*^403/+^ iPSC-CMs, we measured the spatial dispersion of repolarization to examine whether intrinsic differences in repolarization properties of cardiomyocytes in the syncytium become manifested, potentially establishing spatial voltage gradients that might act as a substrate for re-entrant arrhythmia. Field potential durations (FPDs) were measured at each individual electrode in the array (interelectrode distance approximately 300 μm) (Figure 5Ai) with a greater range of repolarization times measured in monolayers of *MYH7*^403/+^ cardiomyocytes compared to *MYH7*^+/+^ (Figure 5Aii). The spread of field potential durations across the 16 electrodes in each array, measured from 10 individual electrode arrays each for *MYH7*^403/+^ and *MYH7*^+/+^ is shown in Figure 5B, demonstrating a consistently greater global dispersion of field potentials in *MYH7*^403/+^ monolayers. Overall, the average standard deviation of FPD within each electrode array increased from 11.4 ± 1.4 ms to 22.5 ± 1.3 ms for *MYH7*^+/+^ and *MYH7*^403/+^ respectively (Figure 5C). Finally, field potential durations were mapped relative to the electrode position in the array showing clear local dispersion of repolarization times of up to 40-50 ms in *MYH7*^403/+^ but not *MYH7*^+/+^ (Figure 5D), potentially resulting in steep voltage gradients that could facilitate re-entrant arrhythmia.

**Figure 4.**
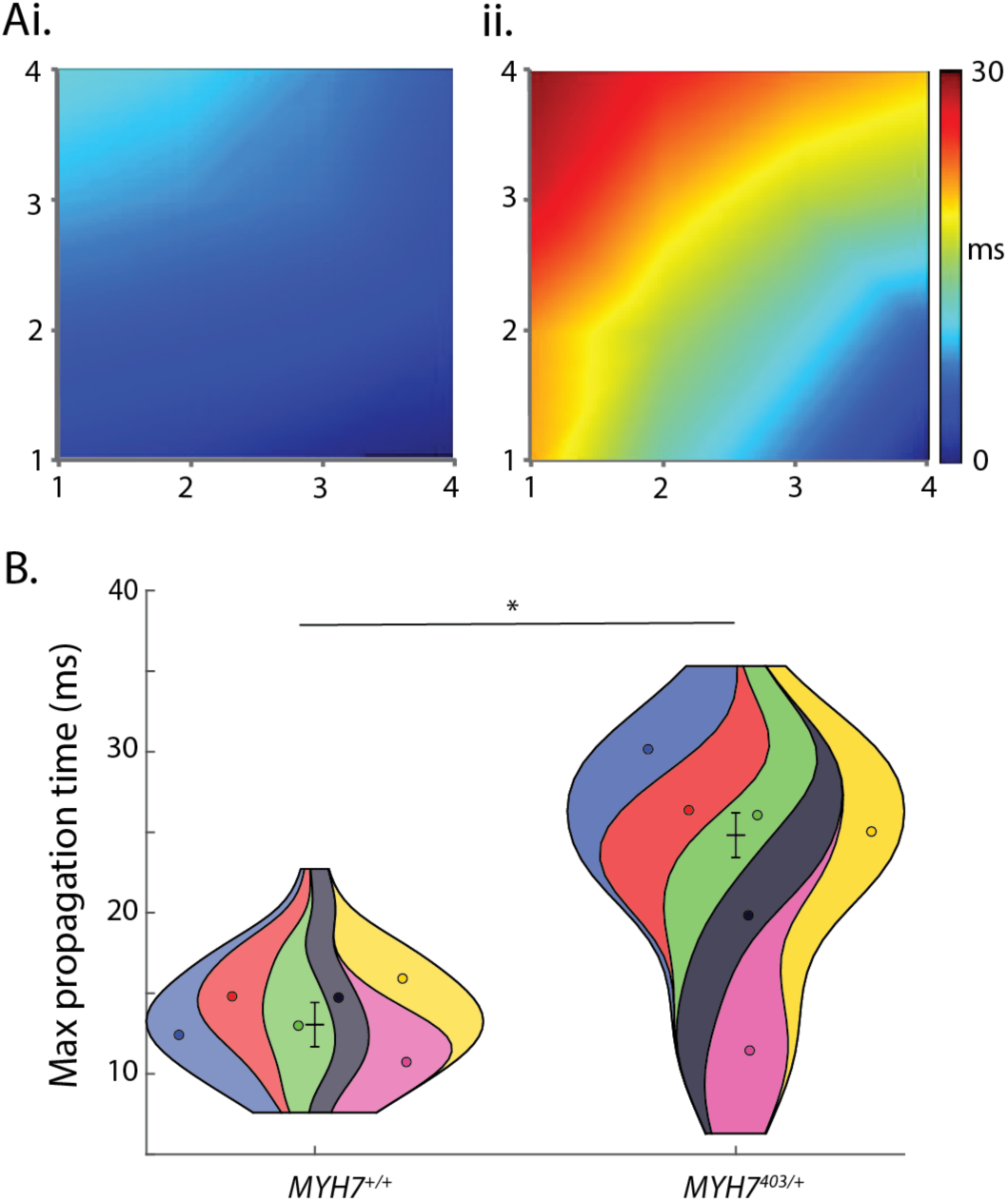
MYH7^403/+^ slows conduction velocity in mono layers of iPSC-CMs. A) Conduction maps for *MYH7*^+/+^ (i) and *MYH7*^403/+^ (ij) colored according to activation time measured on a microelectrode array. B) Violin plots summarizing maximum propagation for *MYH7*^+/+^ (n = 64 replicates/wells from N = 6 differentiations) and *MYH7*^403/+^ (n = 66 replicates/wells from N = 6 differentiations) showing significantly slower conduction in *MYH7*^403/+^.

**Figure 5.**
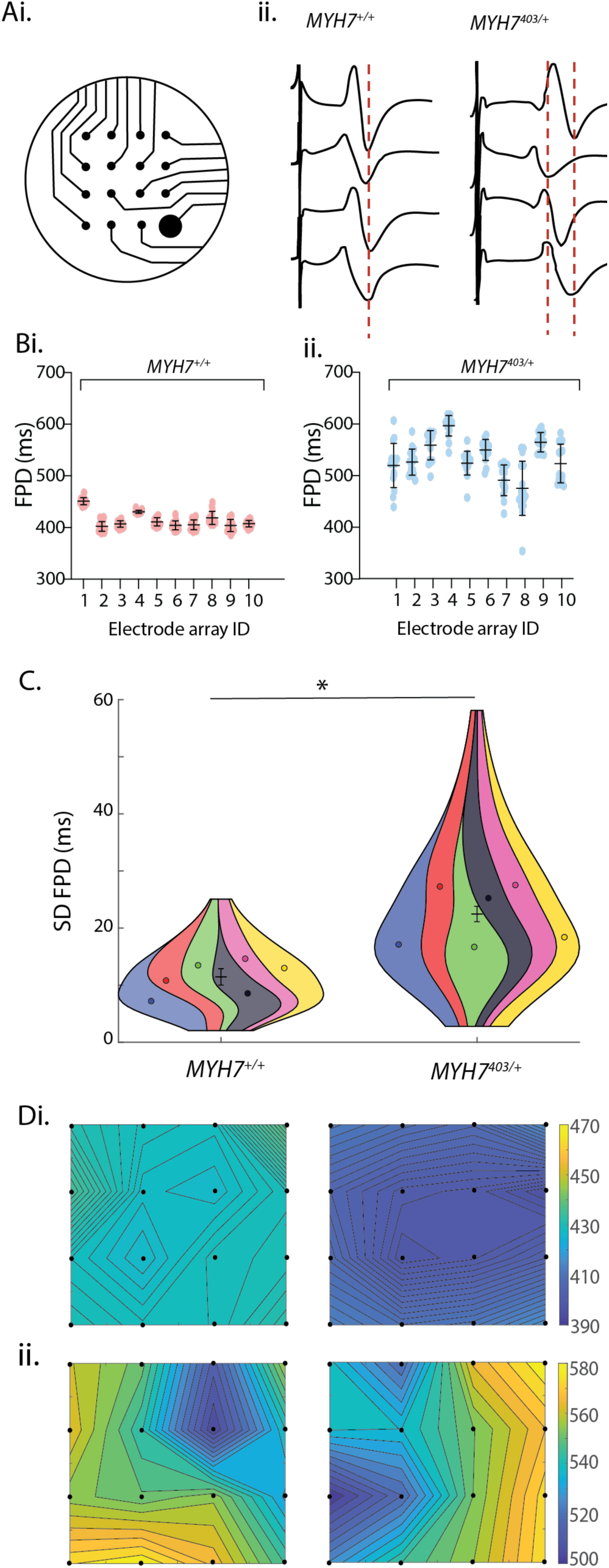
Increased spatial dispersion of repolarization in MYH7^403/+^. Ai) Microelectrode array geometry. ii) Typical field potentials measured in an individual array for *MYH7^+/+^* and *MYH7*^403/+^. B) Range of field potential durations across the 16 electrodes in 10 individual arrays for both *MYH7*^+/+^ (i) and *MYH7*^403/+^ (ij). For each array, individual datapoints represent the field potential duration recorded at an individual electrode in the array. C) Violin plots summarizing the standard deviation of field potential duration in each electrode array for *MYH7*^+/+^ (n = 77 wells from N = 6 differentiations) and MYH7^403/+^ (n = 111 wells from N = 6 differentiations) showing significantly more dispersion of repolarization ti mes for *MYH7*^403/+^. D) Spatial maps of repolarization duration for *MYH7*^+/+^ (i) and *MYH7*^403/+^ (ij).

### 3.5. Molecular basis of electrophysiological phenotype in R403Q

To investigate the molecular basis of the observed differences in electrical phenotype, we measured changes in mRNA and protein expression. First, for mRNA we used a curated panel (nanoString nCounter) targeting cardiac ion channels, calcium-handling proteins and transcription factors. Our data show transcriptional down-regulation of rhythmonome genes involved in calcium handling (*PLN, RYR2*, *SLC8A1*) and cardiac repolarisation (*KCNH2*); sarcomere genes *(MYL7, TTN, TNI1, TNI3);* and genes involved in ventricular cell fate *(NKX2-5, CORIN)* (Figure 6 A/B). to complement this, we finally measured the of rhythmonome proteins using western blot (Figure 7A), showing reduced expression of key proteins involved in cardiac excitation and conduction including 80 ± 12 %, 75 ± 5 %, and 40 ± 10 % reduction in Nav1.5 (the cardiac sodium channel), connexin-43, and Kir2.1 (inward rectifier potassium channel) respectively (Figure 7B).

**Figure 6.**
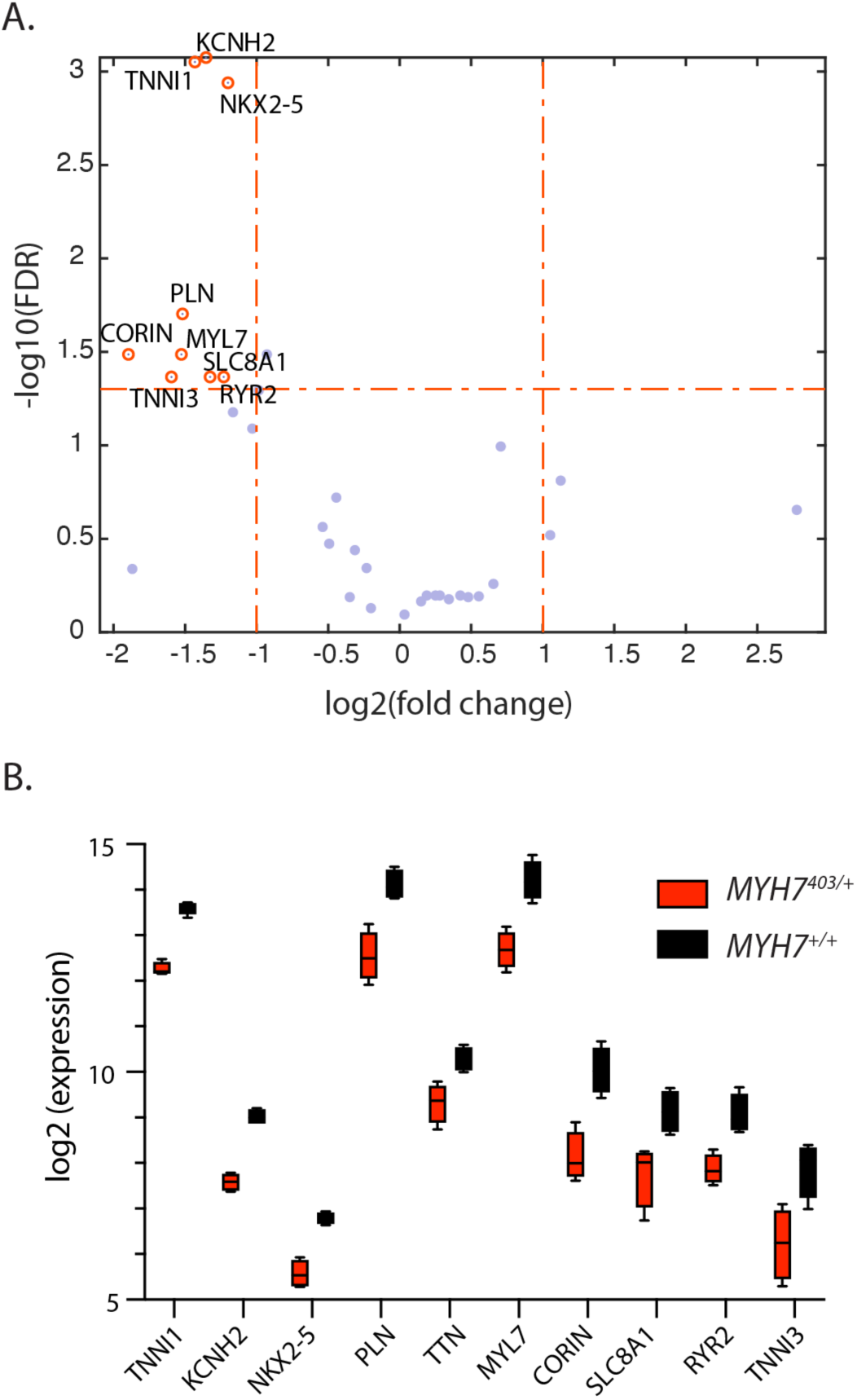
MYH7^403/+^ alters gene expression in iPSC cardiomyocytes. A) Volcano plot showing differentially expressed genes between *MYH7*^+/+^ and *MYH7*^403/+^. B) Comparison of expression of differentially expressed genes.

**Figure 7.**
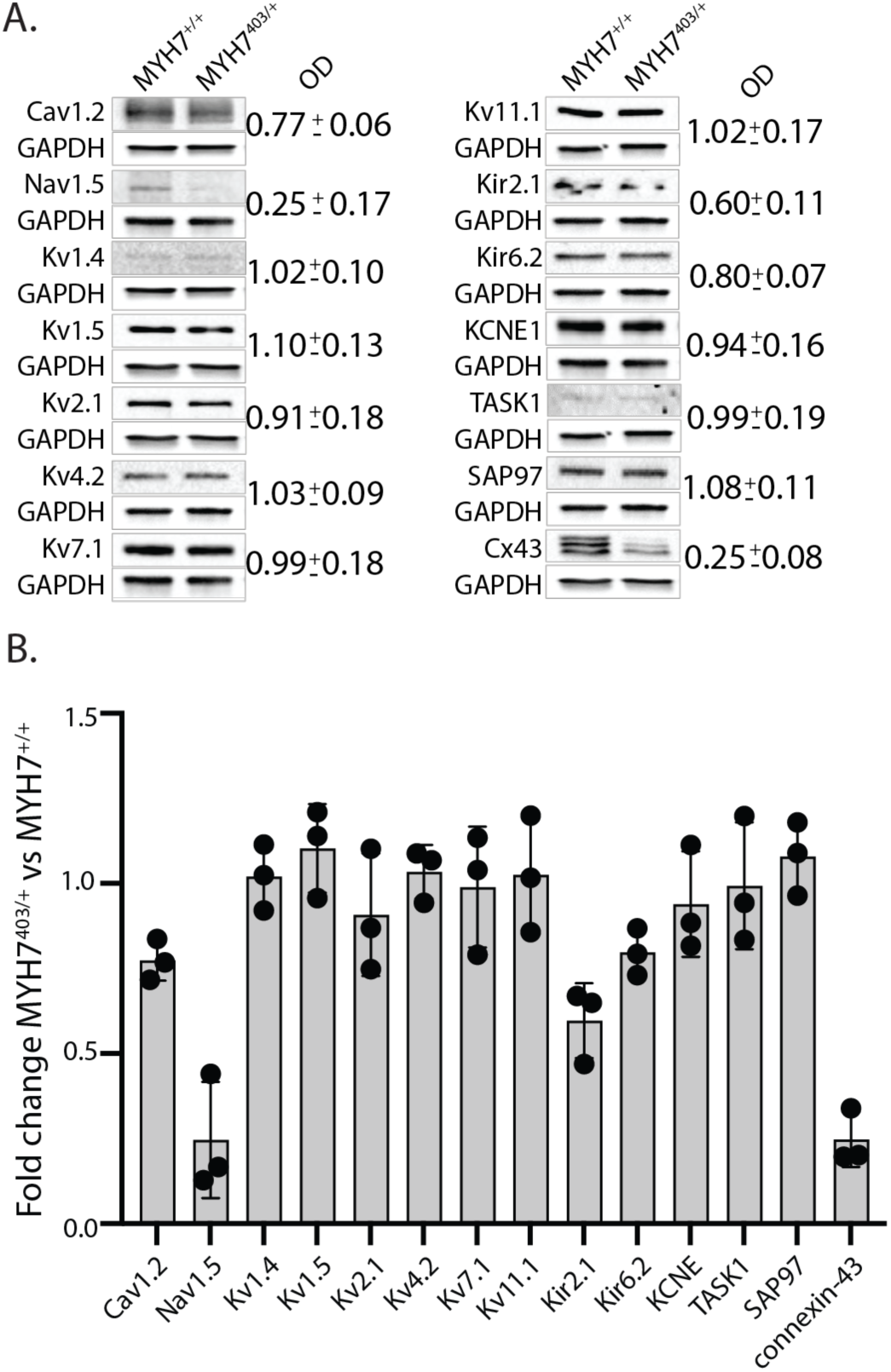
MYH7^403/+^ reduces expression of key proteins involved in cardiac conduction. A) Typical western blots of rhythmonome proteins for *MYH7*^+/+^ and *MYH7*^403/+^. Full Western blots, including replicates, are provided in Supplementary data. B) Summary data showing fold change in expresison (measured via optical densiometry) for rhythmonome proteins.

## 4. Discussion

HCM is an inherited cardiac disorder that results in hypertrophy, fibrosis, myofibre disarray, arrhythmias, and sudden cardiac death^21^. Conduction delays and dispersion of repolarisation are clinically associated with HCM^8,22^. However, the correlation between conduction defects and the histopathology of HCM is poor, meaning the mechanism underlying these electrical phenotypes is unclear. In this study, we used an iPSC model of the *MYH7* R403Q variant to show that reduced conduction velocity occurs due to changes intrinsic to property of *MYH7*^403/+^ cardiomyocytes that occurs due to a dramatic reduction in expression of connexin-43 and sodium channel proteins - both key molecular mediators of conduction in the myocardium. Furthermore, we show that this reduced electrical coupling results in significant spatial dispersion of repolarisation – a well-established proarrhythmic substrate^23,24^ – that may provide a biophysical basis that contributes to sudden arrhythmic death in HCM patients.

### MYH7 R403Q slows conduction in iPSC-derived cardiomyocyte monolayers

HCM is a disease that has been modelled in a range of systems including intact heart muscle strips, isolated cardiomyocytes, myofibrils, purified actomyosin, and animal models^25^. More recently, iPSC-derived cardiomyocyte models of HCM related to variants in *MYH7*^26,27^, as well as other genes^28,29^, have been shown to reproduce key elements of the HCM phenotype *in vitro*, including cellular morphology/hypertrophy^28^, elevated metabolism^30^, disrupted calcium handling, and hypercontractility^26,29^. In relation to electrophysiology, data from iPSC models is less extensive and inconsistent. Studies of *MYH7-*related HCM models in iPSC-derived cardiomyocytes report variable electrical phenotypes, with both unchanged or prolonged action potential duration reported, as well as irregular beating^26,31^. However, there are no *in vitro* studies employing iPSC models that examine macroscopic, ‘tissue level’ electrophysiological properties that might provide a biophysical explanation for the conduction abnormalities seen in patients.

In this study we used microelectrode arrays to record electrical signals from monolayers of iPSC cardiomyocytes, allowing us to study ‘tissue-level’ properties such as conduction velocity and repolarisation dispersion. Our data showed, for the first time, significantly slowed conduction velocity (approximately 50 % reduced;Figure 4) in an *in vitro* model of HCM. Reduced conduction velocity increases the chance of arrhythmia by reducing the spatial scale over which re-entry is possible^32^. In the clinical setting, slow conduction has been reported in patients with HCM^33,34^. Given the histopathology of the HCM myocardium, particularly regarding the presence of interstitial fibrosis, it is generally assumed that disruption of normal electrical propagation is a function of this fibrosis and the associated reduction in coupling between cardiomyocytes^35,36^. However, several studies have suggested that this not necessarily the case. For example, in the clinical setting, Aryana *et al.* reported a lack of concordance between imaging of intramural fibrosis and low voltage on endocardial or epicardial mapping^9^. Similarly, in *MYH7* R403Q in mice, Wolf *et al.* demonstrated that neither the extent nor the location of fibrosis correlated with electrical mapping of conduction properties^10^, nor did this correlate with the propensity for arrhythmia. Furthermore, in a separate mouse study, Hueneke *et al* showed that tachyarrhythmias are observed at a far earlier age than the onset of hypertrophy, suggesting an alternative pathway for arrhythmogenesis in HCM, that is independent of alterations to the structure of the myocardium^11^. Our data are consistent with these observations and support a biophysical basis for conduction slowing in *MYH7* R403Q, intrinsic to cardiomyocyte electrophysiology, that may be accentuated by myocyte disarray in addition to fibrosis and later in the progression of disease.

### Reduced electrical coupling between MYH7 R403Q cardiomyocytes results in increased spatial dispersion of repolarisation

Spatial dispersion or regional differences in repolarisation times facilitate the initiation and maintenance of re-entrant arrhythmias^37^. This view is supported by studies showing that regional differences in repolarisation time lead to an increased susceptibility to arrhythmias in response to premature stimulation^38–40^. Similarly, previous work has shown a tight correlation between the range and variability in action potential duration across the tissue and the duration of the vulnerable window for initiation of re-entry^39^, while on patient ECGs, measurements of T wave area, a metric that reflects dispersion of repolarisation, is predictive of ventricular arrhythmias^41^. Such spatial dispersion of repolarisation is thought to arise because of variation in ion channel gene expression, or ion channel function, in different regions of the heart. However, in normal healthy tissue, tight electrotonic coupling ensures that repolarisation of individual cells, or different regions of the tissue, is synchronised and hence is less vulnerable to arrhythmia^42,43^.

In our iPSC model of *MYH7* R403Q, we hypothesised that the observed reduction in conduction velocity reflected reduced electrical coupling, and that this might alter the degree to which repolarisation was synchronised between cells, manifesting in increased spatial dispersion of repolarisation. This was indeed the case, with significantly greater variability and range of field potential durations (reflective of repolarisation time) measured across R403Q monolayers compared to their CRISPR-corrected controls (Figure 5). Moreover, these voltage gradients occurred over small, local spatial scales. Differences of up to 150 ms in field potential duration were recorded across electrode arrays spaced over areas just 1-2 mm across, with steep voltage gradients occurring even between neighbouring electrodes (see example Figure 5Dii). In relation to arrhythmogenesis, this is important since it is thought that local repolarisation dispersion (as opposed to global differences) often serve as sources for the ectopic beats that trigger arrhythmias ^44,45^. Such dispersion of repolarisation has been linked to arrhythmogenesis in a range of inherited and acquired disorders including long QT syndrome^23,46^, Brugada syndrome^23^, heart failure^47^, and post-ischaemia^48^, where voltage gradients occur variously due to factors including regional differences in ion channel expression as well as electrical uncoupling following gap junction downregulation. Specifically in relation to HCM, Saumarez *et al.* reported fractionated conduction related to myocardial disarray and fibrosis on patient electrograms^49^, while Hueneke *et al.* reported dispersion of repolarisation on a global scale, related to differential expression of potassium channel genes in different regions of the mouse heart^11^. Our observations of highly discordant local repolarisation in R403Q monolayers is therefore consistent with a mechanism that could contribute to arrhythmic susceptibility in patients. To our knowledge, this is the first report of such pronounced local spatial dispersion of repolarisation in a model of HCM, that results solely from the biophysical properties of the HCM cardiomyocyte.

### Molecular basis of electrophysiological changes in R403Q cardiomyocytes

Conduction of electrical signals in the ventricular myocardium is regulated primarily by voltage-gated sodium channels (Nav1.5), which are the main contributor to cardiomyocyte depolarisation, and gap junctions (primarily Connexin 43 in the ventricle) that allow the flow of current between coupled cells. Loss of either of these critical contributors to cardiac conduction can disrupt regular electrical propagation and increase the risk of arrhythmias. Our analysis of protein expression showed a dramatic (∼80%) reduction in the expression of both connexin-43 and Nav1.5 proteins in *MYH7*^403/+^ cardiomyocytes compared to their CRISPR-corrected controls. In addition to this, we also measured a ∼ 50% reduction in Kir2.1 protein (an inward rectifier potassium channel). Functionally, this reduction in Kir2.1 would serve to depolarize the resting membrane potential, leading to functional inactivation of the remaining Nav1.5 population, so acting as a potential ‘third hit’ on electrical conduction.

The degree of reduction in conduction velocity observed (Figure 4), is consistent with previous studies in the literature that have examined a similar loss of connexin expression in other systems. In a mouse model of conditional deletion of Cx43, van Rijen *et al.* showed that a 70 – 95% conditional deletion of Cx43 was necessary to reduce conduction velocity/increase the dispersion of conduction, while a 50 % loss of function had no effect^50^. Similarly, an earlier study by Reaume *et al.* showed conduction slowing of between 42% and 56% associated with a 95% reduction in Cx43 protein expression^51^. As a result of these observations, while reduced expression and/or localisation of Cx43 has been observed in several pathological states, including post-infarction^52^ and in HCM^53^, it has been considered unlikely that the degree of reduced expression reported for these diseases was sufficient to directly affect conduction, and hence promote arrhythmogenesis. Rather, a second factor such as fibrosis or collagen deposition was likely responsible for the increased propensity for arrhythmias in these patients. The mechanism of conduction slowing reported here for R403Q cardiomyocytes is therefore unique in that it is the first time reduced expression of connexin protein has been measured in a model of a HCM gene variant of a sufficient magnitude to directly impact cardiac conduction. Moreover, this effect is likely amplified by a simultaneous reduction in the expression of sodium channels and inward rectifier channel protein in a triumvirate of altered rhythmonome protein expression that coalesce to slow cardiac conduction.

### Transcriptomic analysis of R403Q cardiomyocytes

Transcriptome data showed genes related to calcium handling, cellular electrical repolarisation, sarcomere structure, and cardiac cell fate are differentially expressed between *MYH7*-C^+/+^ and *MYH7*^403/+^ cardiomyocytes. First, expression of *NKX2-5* was 2.3-fold higher in normal cardiomyocytes compared to *MYH7*^403/+^. NKX2-5 is a key transcription factor involved in cardiac development and cell fate across species^54^. Of relevance to this study, NKX2-5 dysregulation is associated with defects in cardiac conduction and electrophysiology^55^, including reduced expression of connexins^56,57^ as well as Nav1.5, RyR2 and Kv11.1^57^, consistent with our transcriptomic and protein data (Figures 6/7). Similarly, a mouse model employing an inducible system for disruption of notch signalling, that resulted in downregulation of NKX2-5, displayed a strikingly similar electrical phenotype to that seen here including slow conduction velocity and irregular beating related to reduced expression of gap junctions and sodium channels^58^. Another gene that was downregulated, which is a target for NKX2-5, was Corin. Corin is a serine protease highly expressed in the heart that has conserved binding sequences for NKX2.5 in its 5’-flanking regions^59^. Corin has previously been identified as a cell surface marker for ventricular cell populations^60^ and has been reported as being dysregulated (both down and up-regulated) in hypertrophy and heart failure ^61–64^. Furthermore, reactome pathway analysis also associates Corin with cardiac conduction^65^. Taken together with the altered expression of NKX2-5 in this model of *MYH7*^403/+^, our transcriptome data is therefore consistent with the *in vitro* electrical phenotypes reported here. Furthermore, if maintained during development and into adulthood in patients, these changes would likely contribute to increased risk of arrhythmias and sudden death.

In relation to calcium handling, we observed downregulation of a cluster of genes including *RYR2* (encoding the ryanodine receptor), *SLC8A1* (encoding the sodium-calcium exchanger), and *PLN* (encoding phospholamban). A wealth of previous studies has reported altered calcium homeostasis in models of HCM^13,26,66–68^ and indeed have identified disrupted calcium handling as central to driving HCM pathology^26,68^. In particular, there was altered expression of key calcium handling proteins, including *RYR2*^68^ – in agreement with the transcriptome data we report here.

The third cluster of genes downregulated in *MYH7*^403/+^ cardiomyocytes was related to sarcomere structure – specifically troponin I (both *TNNI1* and 3 isoforms), myosin light chain 7 (*MYL7*), and titin (*TNN*). Whilst there were no changes in *TNNI1:TNNI3* or *MYL2:MYL7* ratios, that might reflect a less mature or less ventricular cell identity of *MYH7*^403/+^ cardiomyocytes (Supplementary Figure 1), the broad downregulation of sarcomere genes observed here might reflect overall reduced structure and order of the sarcomere associated with this variant in a thick filament protein (myosin heavy chain).

Finally, related to cardiac repolarisation, expression of the *KCNH2* gene, encoding the pore-forming subunit of the rapid delayed rectifier potassium current, one of the main drivers of action potential repolarisation in human ventricles, was reduced. This is consistent with studies in mice, from our group and others, that have shown downregulation of potassium channel gene expression associated with HCM variants in both the *MYH7*^11,20^ and *MYBPC3*^69^. However, in these studies, a reduction in protein expression, or ion channel current density, was also reported in association with the reduced gene expression. Conversely, we see no significant change in protein levels of the ion channel encoded by this gene (Kv11.1) in R403Q vs CRISPR-corrected controls (as well as only modest changes in repolarisation time). Similar results were also reported by Flenner *et al*, where data from either human iPSCs, or from human ventricular samples, showed no reduction in K^+^ currents, in contrast to their mouse data^69^. Such discordance between mRNA and protein levels is common^70^, with specific examples reported regarding potassium channels and arrhythmias^71^, and most likely reflects that protein abundance is in large part controlled at the post-transcriptional and translational levels and that these processes can adapt to compensate for changes in mRNA transcription to keep cellular protein levels at appropriate levels for normal function.

## Conclusions

In this study, we have shown that the R403Q pathogenic variant in *MYH7*, a common and severe cause of HCM, results in changes in cardiomyocyte electrophysiology, that may contribute to proarrhythmia and increased risk of sudden death in patients. Specifically, in monolayers of *MYH7*^403/+^cardiomyocytes, reduced electrical coupling resulted in significantly slowed conduction velocity, and an accompanying increase in spatial dispersion of repolarization that established steep voltage gradients in monolayers of iPSC-derived ‘pseudo tissue’. Analysis of rhythmonome proteins revealed that reduced electrical coupling resulted from reduced expression of the key proteins – connexin-43, Nav1.5 and Kir2.1 - that support electrical conduction between cardiac cells. This is the first report of a cardiomyocyte-intrinsic mechanism of ‘tissue-level’ proarrhythmia in a model of HCM that may represent a new focus for targeted antiarrhythmics in these patients. As such it represents a novel, biophysical basis for proarrhythmia in HCM that may be accentuated by structural changes in the myocardium later in the progression of disease to contribute to sudden arrhythmic death in these patients.

## Limitations

Whilst we observe proarrhythmic electrical phenotypes (slow conduction, irregular beating, increased dispersion of repolarisation) in our iPSC model of R403Q we did not observe overt re-entrant activity in our experiments. A likely explanation for this is the small spatial scale of the monolayers studied (microelectrode array geometry is on the order of 2 mm) that, even in the context of slow conduction and discordant repolarisation, were not sufficiently large to support reentry. It may also be the case that changes in the expression of rhythmonome proteins observed here, and their associated phenotypes, may be accentuated by the relatively immature properties of iPSC cardiomyocytes and may be less prominent in more mature cells. Finally, data from our study was acquired from a cell line from a single patient (and it’s CRISPR-corrected isogenic control). Further studies including cell lines from multiple patients with the same variant in *MYH7* would help quantify the role of the genetic background in fine-tuning the emergent electrical phenotype of the R403Q variant.

## Acknowledgements

We thank the Victor Chang Innovation Centre, funded by the New South Wales Government.

